# Scooby-2: fine-tuning AlphaGenome for single-cell genomic track prediction

**DOI:** 10.64898/2026.10.28.762238

**Authors:** Johannes Hingerl, Julien Gagneur

**Affiliations:** School of Computation, Information and Technology, Technical University of Munich, Munich, Germany; Munich Center for Machine Learning, Munich, Germany; Institute of Human Genetics, School of Medicine, Technical University of Munich, Munich, Germany; Computational Health Center, Helmholtz Center Munich, Neuherberg, Germany

## Abstract

**Motivation:** Sequence-to-function models predict genomic tracks from DNA sequence. AlphaGenome has been shown to outperform previous models, including Borzoi, on most track-prediction and variant-effect benchmarks. However, like its predecessors, it predicts tracks only for predefined, mostly bulk-derived cell types and tissues. While Scooby has extended Borzoi to single-cell resolution, a single-cell adaptation of AlphaGenome is still lacking.

**Results:** We present Scooby-2, which combines AlphaGenome with a cell-specific decoder to predict single-cell RNA-seq coverage and chromatin-accessibility genomic tracks from DNA sequence. The package provides the implementation, the parameter-efficient fine-tuning setup, the training pipeline and inference scripts. When evaluated on the NeurIPS bone marrow single-cell dataset, Scooby-2 achieved a Pearson correlation of 0.56 when predicting cell-type-specific gene expression differences, compared with 0.53 for Scooby. When using the distilled AlphaGenome backbone, variant effect prediction concordance of fine-mapped eQTLs from the OneK1K cohort improved from 0.37 to 0.47 Spearman’s rank correlation across all associations, an improvement consistent with that of AlphaGenome over Borzoi itself.

**Availability and implementation:** Scooby-2 alongside training code is available at github.com/gagneurlab/scooby under the MIT license. Trained weights (huggingface.co/johahi/neurips-scooby-2, huggingface.co/johahi/onek1k-scooby-2) are derivatives of AlphaGenome and are distributed under the AlphaGenome Model Parameters Terms of Use.

## Introduction

Sequence-to-function models are trained to predict genomic tracks, i.e., omics measurements with values along the genome, directly from genomic sequence and provide a framework for studying cis-regulatory activity and genetic variation. Enformer (Avsec *et al*. 2021) and Borzoi (Linder *et al*. 2025) established long-range sequence-based models of chromatin and transcriptional tracks, while AlphaGenome extended this paradigm to a broader set of genomic tracks at up to base-pair resolution (Avsec *et al*. 2026). AlphaGenome was reported to improve on external models (including Borzoi) across most genomic-track and variant-effect prediction tasks. AlphaGenome nevertheless produces a fixed collection of tracks learned predominantly from bulk-derived assays. Because each track requires its own output, this design cannot scale to datasets with potentially millions of individual cells.

To generalize any bulk sequence-to-function model to single-cell omics, we previously introduced Scooby (Hingerl *et al*. 2025). Scooby combines Borzoi’s sequence representation with a lightweight decoder whose parameters are generated from a low-dimensional embedding of each cell, allowing predictions for any cell. The resulting model predicts strand-specific scRNA-seq coverage and scATAC-seq insertion tracks. Scooby was shown to recapitulate cell-specific expression of held-out genes and resolve cell-type-specific regulatory effects.

Here, we extend Scooby to the AlphaGenome backbone. Together with a configuration-driven training and inference interface, we provide an AlphaGenome-based framework for single-cell genomic track prediction.

## Implementation

For AlphaGenome, we leveraged the AlphaGenome-Pytorch implementation (Bredikhin *et al*. 2026). For computational efficiency, we opted to keep the model input at 524,288 bp, as this was shown to not materially affect gene expression prediction or variant effect prediction (Avsec *et al*. 2026, Figure 7). We considered a 1-bp as well as a 32-bp resolution output matching the AlphaGenome native resolution and the original Scooby resolution, respectively. A multilayer perceptron maps each low-dimensional single-cell embedding to a cell-specific convolution that predicts strand-specific RNA coverage and ATAC insertion tracks along the genome (Fig. 1a).

**Figure 1.**
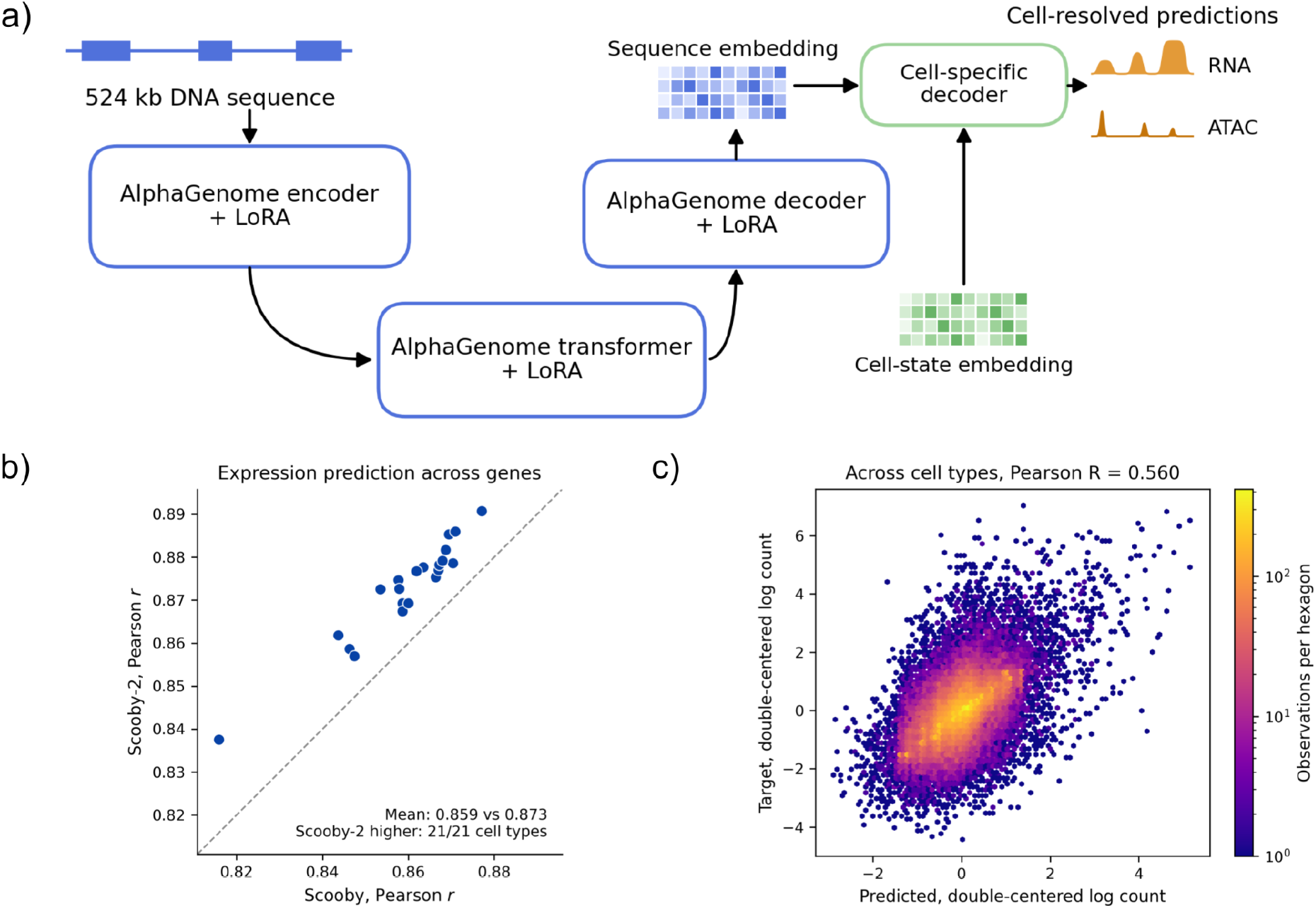
**a)** Scooby-2 architecture, including the AlphaGenome backbone and the cell-specific decoder. **b)** Held-out expression correlation for Scooby-2 on the NeurIPS dataset against Scooby across genes. Each point is a cell type. **c)** Predicted against measured between-cell-type deviations of gene expression for Scooby-2 on the NeurIPS dataset.

For the 32-bp model, we truncated the AlphaGenome decoder after its second upsampling block, where it reaches 32-bp resolution. We replaced the higher-resolution path with a freshly initialized 32-bp output embedder. As in Borzoi-based Scooby, predictions were center-cropped to 6,144 bins (196,608 bp). We refer to this model as Scooby-2.

The sequence backbone was fine-tuned using low-rank adapters (LoRA, Hu *et al*. 2021), while the cell-specific decoder and 32-bp output embedder were trained directly when applicable. Furthermore, an optional GroupNorm layer on the sequence embedding was used to stabilize training for scRNA-seq data (Wu and He 2018). For compatibility with AlphaGenome’s weight-standardized convolutions (Brock, De, and Smith 2021), convolutional low-rank updates were normalized jointly with their pretrained filters. Training used reverse-complement and shift augmentation together with the Poisson-multinomial loss from Linder *et al*. (2025). A run is described by one YAML file containing paths and relevant hyperparameters. Paths are resolved relative to the config file and checked before the model is instantiated. Scooby can then be loaded with:

~~~
from scooby import ScoobyAlphaGenome
model = ScoobyAlphaGenome.from_pretrained(“checkpoint/”, device=“cuda”)
~~~

## Results

We evaluated Scooby-2 on the NeurIPS 2021 bone-marrow multiome dataset used to benchmark Scooby (Luecken *et al*. 2021). To enable a direct comparison, Scooby-2 and Scooby used the same genomic held-out folds (AlphaGenome ‘fold1’), single-cell targets and Poisson-MultiVI cell-state representation (Martens *et al*. 2024). Genes overlapping the validation and test regions were excluded when constructing the single-cell embedding, preventing the information from held-out genes from entering through the cell embedding.

On genes overlapping the test set, Scooby-2 achieved a mean Pearson correlation of 0.87 between predicted and observed expression across genes within cell types, compared with 0.86 for Scooby (Fig. 1b). To assess whether the models captured relative differences between cell types rather than gene abundance alone, we additionally evaluated the double-centered gene-by-cell-type expression matrix after removing gene and cell-type means as in Hingerl *et al*. (2025). Here, Scooby-2 reached a Pearson correlation of 0.56, surpassing Scooby by 0.03 (Fig. 1c).

We next tested whether adapting AlphaGenome’s convolutional encoder improved transfer to single-cell data. For this, we trained an otherwise identical model without convolutional adapters. This ablation retained LoRA of the transformer attention and feed-forward layers and trained the same 32-bp output embedder and cell-specific decoder. Including the convolutional low-rank adaptation improved across-gene correlation from 0.86 to 0.87 and cell-type-specific correlation from 0.55 to 0.56. The increase in cell-state-specific gene expression prediction might justify the computational cost (∼1.25x training time) of adapting the convolutional encoder for this task.

We also tested a base-pair-resolution version using the full AlphaGenome decoder, which did not improve expression prediction.

Finally, we evaluated variant-effect prediction for fine-mapped eQTLs in peripheral blood cell types. As in Avsec *et al*. (2026), we scored variants with a model initialized from the AlphaGenome all-folds weights (the distilled student), which we fine-tuned on the OneK1K single-cell resource (Yazar *et al*. 2022) using all genomic folds. Both AlphaGenome pretraining and Scooby-2 fine-tuning were done using the entire genome, leaving no held-out genes for evaluation. As in the AlphaGenome study, only the generalization capacity of the model at predicting variant effects was considered. Training was only stable when the sequence embedding was normalized before the cell-specific decoder by adding a GroupNorm, otherwise, predictions collapsed towards zero early in training. As a baseline, we used the Scooby model trained on the same OneK1K data.

We selected OneK1K associations with a posterior inclusion probability above 0.9 from the eQTL Catalogue (Kerimov *et al*. 2021), giving 1,310 associations (762 variant-gene pairs) across 21 cell types after excluding variants outside the 524 kb input window. For each association, we predicted the expression of the gene using the sequences of the reference and of the alternative alleles. We summed the predicted counts over the gene body and over all cells of the associated cell type, and averaged over forward and reverse-complemented inputs, as we previously described (Hingerl *et al*. 2025). The resulting log_2_ fold change was compared with the eQTL effect size (beta). Scooby-2 increased the Spearman correlation with the effect size from 0.37 to 0.47, and exceeded Scooby in 8 of the 10 cell types with more than 20 associations (Fig. 2a). When faceted by distance from the TSS, both models showed diminishing performance for distal variants (Fig. 2b). Nonetheless, Scooby-2’s gain over Scooby was largest for variants more than 35 kb from the transcription start site. Here, Scooby-2 predicted the direction of effect with a cell-type-weighted sign AUROC of 0.74, compared with 0.64 for Scooby, and reached a Spearman correlation of 0.35, compared with 0.20 for Scooby. Altogether, these results are consistent with AlphaGenome’s improvement over Borzoi in eQTL effect prediction (Avsec *et al*. 2026).

**Figure 2.**
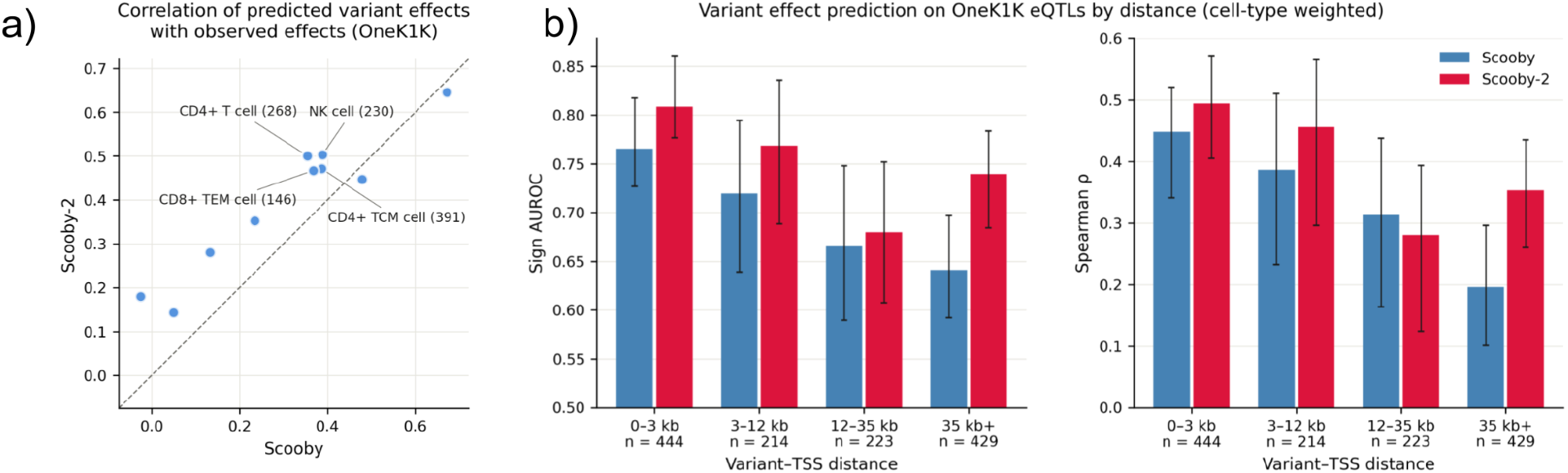
**a)** Spearman correlation of predicted effects (log_2_ fold change) with observed eQTL effect size (beta) for Scooby-2 against Scooby. Each point indicates a cell type from the OneK1K cohort. Only cell types with more than 20 associations are shown. Paired Wilcoxon test *P* = 0.01 across cell types. **b)** Distance-stratified evaluation of Scooby-2 (red) and Scooby (blue) on AUROC of predicted sign vs. observed sign, computed per cell type and averaged weighted by number of associations (left) and Spearman ρ (right). Error bars show 95% bootstrap confidence intervals.

## Conclusion

Scooby-2 adapts Scooby’s efficient approach to sequence-based modeling of single-cell multiome data to AlphaGenome. On a bone-marrow multiome dataset, Scooby-2 exhibited improved prediction of cell-type-specific expression differences. On a single-cell RNA-seq dataset with cell-type-specific fine-mapped eQTLs, Scooby-2 demonstrated improved variant effect prediction over Scooby, while nonetheless still being prone to the decay in variant effect prediction fidelity reported for sequence-to-function models (Karollus, Mauermeier, and Gagneur 2023). Altogether, this work further demonstrates the generality of the Scooby architecture and unlocks AlphaGenome’s potential for single-cell regulatory genomics.

## Acknowledgements

We thank Aleksandr Neverov, Mariia Minaeva, Laura Martens and Nils Wagner for the helpful discussions and feedback. J.G. acknowledges support from the Deutsche Forschungsgemeinschaft (SFB/TransRegio TRR267, Project-ID 403584255) and J.H. and J.G. from the European Research Council (ERC) (EPIC, Grant number: 101118521). The IT Infrastructure of the Chair of Computational Molecular Medicine was funded by the DFG under the project numbers 553375143 and 461264291. AI assistance was used in the preparation of this work. Anthropic’s Claude was employed for text editing and proofreading, as well as for code generation. The authors take full responsibility for all content presented in this manuscript.

